# Assessment of EMS-induced mutagenesis in *Fagopyrum esculentum* Moench (Buckwheat)

**DOI:** 10.64898/2026.04.25.720850

**Authors:** Ayesha Badar, Iram Siddique, Hakeem Mubeen

## Abstract

Global demand for pseudocereals, including buckwheat, has surged in recent years due to their higher nutritional and pharmaceutical value than cereals and also due to them being a climate-resilient, gluten-free, and potential crop for combating cancer, type ll diabetes, and overcoming micronutrients hidden hunger problems that lack in cereals. Major efforts are needed to make its cultivation more popular by improving its quantitative and qualitative traits through crop genetics by adopting modern genetic, molecular, and mutational approaches, which also necessitate the induction of genetic variation for better yielding and improved varieties. In this experimental study, the induced mutant populations of widely recommended VL-7 and PRB-1 varieties of buckwheat were generated using different concentrations treatments of ethyl methane sulfonate (EMS). Investigation on induced phenotypical and genotypical variations in individual plants of M_1_ population of different treatments resulted in morphological and cytological mutant types affecting plant germination, survival, height and morphology, leaf morphology, flower morphology, growth period, chlorophyll and pigments abnormalities in leaves, leaf growth pattern, plant fertility, yield, and cytological aberrations. This experiment showed that plant survival decreased with the concentration of the mutagen doses. The lower doses resulted in dwarf varieties suitable for cultivation as they increased yield by having higher breaking force and lower lodging index over the tall plants. Studies on various quantitative parameters revealed the general effectiveness of intermediate doses and stimulatory effectiveness of lower and higher concentrations in M_1_ generation.

## Introduction

Global food security relies heavily on a few major cereals like wheat, rice, and corn, which provide over 50% of global calories to the human diet but often lack essential micronutrients, resulting in deficiencies leading to “hidden hunger” (De Valença *et al*., 2017). To address this, interest has grown in nutrient-rich pseudocereals, which also produce many secondary metabolites that act as an important source of pharmaceutical drugs. Hence, known as “grains of the twenty-first century” for their high nutritional value and health-promoting compounds (FAO, 2011). Buckwheat (*Fagopyrum esculentum* Moench) is a dicotyledonous crop belonging to the family Polygonaceae (2n = 2x = 16) was neglected during the 20th century in western countries because of the increased yield of wheat during the Green Revolution (Cawoy *et al*., 2008). The mineral content in pseudocereals like buckwheat is about twice as high as in other cereals, which find immense potential to address the hidden hunger problem. As the agricultural land is a limiting factor, buckwheat is climate resilient crop and can be grown even on marginal lands, in poor soils, in conditions not fit for cereals, and can withstand such abiotic stresses as drought, temperature, salinity, and heavy metal stress; thus, are considered a future crop to tackle malnutrition and future food crises. It contains a high level of polyunsaturated essential fatty acid such as linoleic acid, vitamin B_1_, E, and C, and is also gluten-free. It could therefore be used as a substitute for wheat in gluten-free diets for celiac patients. Buckwheat seeds have also been shown to exhibit very high anti-oxidant activity, even higher than that of other grains like oats, barley, wheat, rye, etc.

Buckwheat embryos are unique in accumulating galactosyl cyclitols, not raffinose oligosaccharides. Being an excellent phosphorus scavenger, buckwheat is often regarded as a natural phosphorus pump having 10 times higher uptake capacity than wheat (Zhu *et al*., 2002). These are enriched with various bioactive compounds such as polyphenols, flavonoids, amino acids, dietary fibre, antioxidants, and other essential components like fagopyritols. Rutin is the most abundant flavonoid found in buckwheat, which is absent in cereals and other pseudocereals. Rutin has been widely used in the treatment of edema, haemorrhagic diseases, hypertension, and inflammation (Omidbaigi & Mastro, 2004; Nile & Park, 2014). It is also known to avert Alzheimer’s disease by ameliorating oxidative stress (Javed *et al*., 2012). Buckwheat also contains fagopyrin, which is used in photodynamic therapy for the treatment of cancer, diabetic, and microbial cells (Dai *et al*., 2009; Amezqueta *et al*., 2012; Benković *et al*., 2014). Buckwheat has SDF (soluble dietary fibre) that decreases human blood cholesterol level, and its proteins are known to reduce cholesterol concentration in the serum by increasing the faecal excretion of the steroids (Takahama & Hirota, 2011), and they also suppress colon carcinogenesis by reducing cell proliferation (Liu *et al*., 2001). Buckwheat has higher resistant starch and flavonoid contents, providing low glycemic and insulin indexes (Stokić *et al*., 2015; Gao *et al*., 2016), advantageous for diabetic control. Buckwheat has 17 amino acids, out of which 9 are essential. The presence of higher levels of amino acids, especially arginine, aspartate, and lysine than cereals, which are generally considered deficient in essential amino acids, makes buckwheat a promising alternative for super food applications (Janssen *et al*., 2016; Sytar *et al*., 2018). Additionally, it is also considered as a mineral source having high concentrations of copper, magnesium, phosphorus, potassium, and zinc compared to major cereals (Campbell, 1997), along with the presence of well-documented useful vitamins (Alvarez-Jubete *et al*., 2010). Buckwheat improvement in India has not achieved reasonable success so far because of the marginal importance it receives in the national planning and competition from commercial crops. Despite its adaptability and nutritional value, its cultivation has declined due to competition from commercial crops, prompting the Indian Council of Agricultural Research (ICAR) to initiate focused efforts since 1982 under the AICRP on Under-Utilized Crops, releasing three improved varieties—Himpriya, VL-7, and PRB-1—and including it in the Jai Vigyan Project for its food and nutritional potential. Although the genus *Fagopyrum* has a significant amount of genetic diversity in its wild and cultivated forms, it faces some major challenges, such as low and unstable yields, an indeterminate growth habit, seed shattering, lodging, and the low shelf life of its flour. The present study was aimed at enhancing the phenotypic and genotypic mutations in the M_1_ generation of buckwheat following mutagenesis with different concentrations of EMS treatments for achieving desired plant characteristics of agronomic and economic importance.

## Materials and methods

### Agro - climatic conditions of the site of study

The present study was conducted at a site, the Botanical Garden (Aligarh Muslim University) in Aligarh. Aligarh has a semi-arid and sub-tropical climate with hot, dry summers and cold winters. The average rainfall in this district is 847.30 mm, and more than 85% rainfall occurs from mid-July to September. During the summer, the average temperature is 36^0^ C and during the winter, the average temperature is 16^0^ C. The soil in Aligarh is sandy loam and alkaline in nature.

### Biological Material and Experimental Procedures

#### Plant Material

The plant material used in this study consisted of two buckwheat (*Fagopyrum esculentum*) varieties: VL-7 and PRB-1.

#### Mutagenesis Treatment

Chemical mutagenesis was performed using Ethyl Methane Sulfonate (EMS) to induce genetic variability. A fresh aqueous stock solution of EMS was prepared using phosphate buffer at pH 7.0. Working solutions of EMS were prepared at concentrations of 0.01%, 0.02%, 0.03%, and 0.04%.

#### Treatment Method

Dry and healthy seeds were pre-soaked in distilled water for nine hours. After soaking, the seeds were treated with EMS solutions for six hours at room temperature (25±2 °C) with intermittent shaking. Control seeds were soaked only in distilled water. After treatment, seeds were thoroughly rinsed under running tap water for 30 minutes to remove any residual mutagen.

#### Germination and Growth Conditions

For germination analysis, 10 seeds from each treatment group were placed on moist cotton in Petri dishes and incubated in a BOD chamber at 27±1 °C. to determine the percentage of seed germination and root and shoot lengths were measured to assess seedling growth. Additionally, 15 seeds per treatment (including control) were sown in nine-inch earthen pots containing a growth medium composed of farmyard manure, soil, and sand in a 1:1:1 ratio. Seeds were sown one to two centimetres deep, maintaining equal spacing, and the pots were placed in a net house at the Department of Botany, Aligarh Muslim University, Aligarh.

#### Studies in M_1_ Generation

The following parameters were studied from M_1_ Generation:

#### Seed Germination

After recording germination counts, the percentage of seed germination was calculated on the basis of the total number of seeds sown in the petri dishes and in the field.

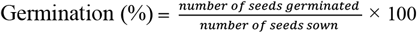

#### Seedling height

By measuring the root and shoot length for each treatment and control, seedling height was recorded after 10 days. Seedling injury was measured in terms of the reduction in seedling height with respect to the control.

#### Plant Survival

The surviving plants in different treatments and control, were counted at maturity, and the survival was computed as percentage of germinated seeds in the field. The following formula was used to calculate the percentage of inhibition injury on reduction.

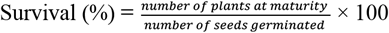

#### Pollen fertility

Pollen fertility was determined from 30 randomly selected plants (10 plants from each replicate) from each treatment and control for both varieties at the time of flowering. Pollen grains were stained with one percent acetocarmine solution on glass slides and covered with cover slips. Pollen grains that took stain and had a regular outline were considered fertile, while the shrunken and unstained ones were considered sterile.

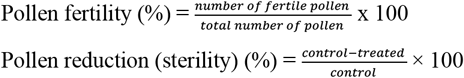

#### Cytological studies

Cytological studies were carried out on the pollen mother cell by fixing younger flower buds from each treatment as well as the control. The purpose of fixation is to kill the tissue without causing any disorder of the components to be studied. It should not only increase the visibility of chromosome structure but should also clarify the details of chromosome morphology and the primary and secondary constriction.

#### Fixation of flower buds

For meiotic studies, young flower buds from 40-45 randomly selected plants were fixed in prepared Carnoy’s fluid (alcohol: chloroform: acetic acid in a 6:3:1 ratio) supplemented with crystals of ferric chloride for 24 hrs. The material was then washed and preserved in 70% alcohol (v/v), then the anthers were squashed in 1% propinocarmine (w/v), dehydrated in NBA series (50% acetic acid+50% normal butyl alcohol), and mounted in canada balsam. More than 600 diving PMCs from each treatment, as well as the control population, were studied and analysed at Metaphase I/II, Anaphase I/II, and Telophase I/II stages. The photomicrographs were taken from temporary as well as permanent slides under the aid of a Nikon photomicrographic unit using a 10X eyepiece and 100X objective lens.

#### Estimation of chlorophyll and carotenoid contents

The estimation of chlorophyll and carotenoid contents was recorded from secondary emerging leaflets of the seedlings raised from mutagen-treated seeds and from control seeds in pots. The procedure of their estimation is elaborated as follows: 100 mg of finely cut fresh leaves was ground to a fine pulp using a mortar and pestle after pouring 10 ml of 100% acetone. The mixture was centrifuged at 5,000 rpm for five minutes at 25 °C. The pellet was discarded, and the supernatant was read using a spectrophotometer. The absorbance in terms of optical density was read at 645 nm for chlorophyll b, 662 nm for chlorophyll a, and 470 nm for carotenoid against the acetone (80%) as blank on a spectrophotometer (Spectronic 20D, Milton Roy, USA). The chlorophyll and carotenoid contents present in the extracts of leaves were calculated according to the equation given by Arnon (1949).

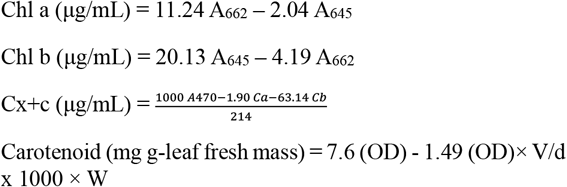

#### Quantitative Traits

The following quantitative traits were thoroughly studied in M_1_ generation:

#### Plant height (cm)

Plant height was measured at maturity in centimetres from the base up to the apex of the plant.

#### Number of fertile branches per plant

The number of fertile branches per plant was calculated at maturity after the completion of flowering in plants.

#### Seed Weight (g)

Seed weight was calculated randomly for 100 seeds of each treatment and control.

#### Total seeds yield per plant (g)

Seed yield per plant was calculated as the weight of the total number of seeds harvested per plant, and the yield of each plant was recorded in grams.

#### Statistical analysis

Data collected for quantitative traits in M_1_ generation were subjected to statistical analysis to assess the extent of induced variation, as indicated below:

#### Mean 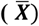

The mean was computed by taking the sum of all the values (X _1_, X_2_………. X n) and dividing by the total number of values (N) involved, thus;

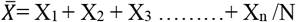

Where, X_1_, X_2_ X n = Observations

N = Total number of observations involved

#### Standard deviation (S.D)

The standard deviation was calculated by the following formula for each parameter of the study.

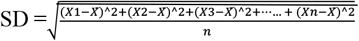

or 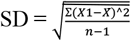

Where, S.D = standard deviation

∑ = sum of all individual aberration is X_1_+ X_2_ +X_3_ + …………X_n_

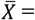 Mean of all observations

N = number of all observations

#### Coefficient of variation (C.V.)

It measures the relative magnitude of variation present in observations relative to their magnitude of mean. It is defined as the ratio of standard deviation to the arithmetic mean expressed as a percentage and a unit-less number. The following formula was applied to compute the coefficient of variability (C.V.)

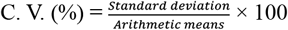

#### Mean Standard error (S.E.)

It is the measure of the uncontrolled variation and can be calculated by the following formula:

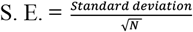

## Results

Mutagenesis is an important method to bring about the variability in plants in a short duration of time. The experiment was undertaken to study the effect of individual treatments of EMS on buckwheat variety VL-7 and PRB-1. Different parameters, like biological and quantitative characters, were studied in this experiment. The results are discussed below.

### Biological study

#### Seed germination and percentage inhibition (In pots)

The germination started on the eighth day after sowing in mutagen-treated plants and in the control of PRB-1. But the germination in control of VL-7 was late as compared to mutagen-treated plants; the germination started almost one week after the germination in mutagen-treated plants. The percentage germination increased with decreasing mutagen concentration in VL-7 and PRB-1. The lowest seed germination was recorded in the case of control of VL-7, which then decreased from 0.01 % EMS concentration to 0.04% EMS. Maximum germination was seen in the control of PRB-1. Percentage inhibition was observed to be increased with increasing concentration of mutagen. Maximum inhibition was observed in 0.04% EMS concentration of both varieties.

#### Seed germination and percentage inhibition (In Petriplates)

Percentage germination and inhibition were also recorded in petriplates under laboratory conditions. Inhibition was found to be increased from 0.01% EMS to 0.04% EMS with a simultaneous decrease in germination in both the varieties (Table 1 & Figure 1).

**Table 1:** Effects of EMS on seed germination, plant survival, and pollen fertility in two varieties, VL - 7 and PRB -1 of *F. esculentum*.

| TREATMENT | GERMINATION (%) | INHIBITION (%) | PLANT SURVIVAL (%) | LETHALITY (%) | POLLEN FERTILITY (%) | REDUCTION (%) |
| --- | --- | --- | --- | --- | --- | --- |
| **VL-7** |  |  |  |  |  |  |
| CONTROL | 26.40 | — | 100.00 | — | 85.71 | — |
| EMS (%) |  |  |  |  |  |  |
| 0.01 | 66.00 | -150.00 | 60.00 | 40.00 | 80.00 | 6.66 |
| 0.02 | 52.80 | -100.00 | 50.00 | 50.00 | 66.60 | 22.29 |
| 0.03 | 52.80 | -100.00 | 50.00 | 50.00 | 100.00 | -16.67 |
| 0.04 | 39.60 | -50.00 | 50.00 | 50.00 | 60.00 | 29.99 |
| **PRB-1** |  |  |  |  |  |  |
| CONTROL | 100.00 | — | 100.00 | — | 88.88 | — |
| EMS (%) |  |  |  |  |  |  |
| 0.01 | 85.60 | 0.144 | 85.71 | 14.29 | 85.71 | 3.50 |
| 0.02 | 72.40 | 0.276 | 100.00 | — | 75.00 | 5.61 |
| 0.03 | 68.00 | 0.320 | 75.00 | 25.00 | 90.00 | -1.26 |
| 0.04 | 51.08 | 0.489 | 80.00 | 20.00 | 83.33 | 6.24 |

**Fig.1(a).**
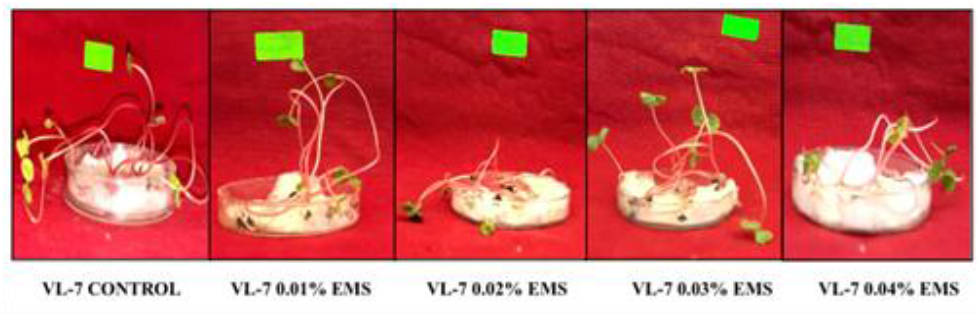
Seed germination pattern in *F. esculentum* variety in VL-7 in control and EMS treatment (0.01% - 0.04%)

**Fig.1(b).**
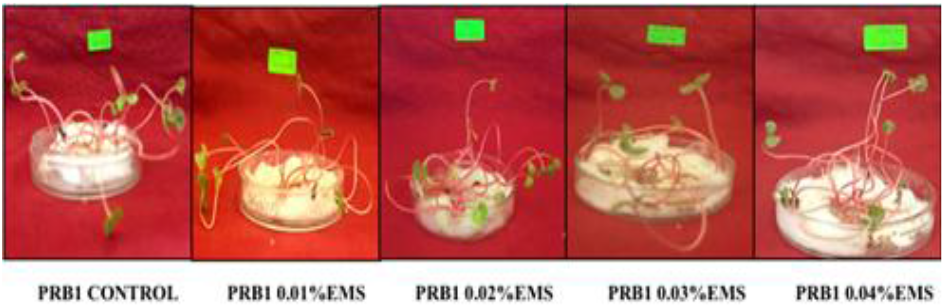
Seed germination pattern in *F. esculentum* variety in PRB-1 in control and EMS treatment (0.01% - 0.04%)

#### Plant survival

The survival of seedling increased with a decrease in the dose of mutagenic treatments. The survival of plants in both the varieties VL-7 and PRB-1 at maturity was 100% in control and then decreased from concentration of 0.01% EMS to 0.04% EMS concentration (Table 1).

#### Pollen fertility

The average pollen fertility in the control set was 85.71% in VL-7 and 88.88% in PRB-1, and then decreased from 0.01% EMS to 0.04% EMS concentration from 80 % to 60% in VL-7 and 85.71% to 83.33% in PRB-1 (Table 1).

#### Morphological studies

In both the varieties VL-7 and PRB-1, the effect of the EMS was studied on various morphological characters, viz., plant height (cm), number of branches per plant, number of fertile branches, bushy character, days to flowering, and days to maturity, weight of the seeds in *F. esculentum* are summarized in the tables below.

#### Shoot length of seedling

The shoot length in control of VL-7 and PRB-1 was 16.15 cm and 15.98 cm, respectively. It decreased from 0.01% EMS to 0.04% EMS in both the varieties from 15.93 cm to 11.50 cm in VL-7 and from 15.60 cm to 13.10 cm in PRB-1, and the mean value of control was 1.83 for both the varieties and decreased from 0.01% EMS to 0.04% EMS from 2.10 cm to 1.00 cm in both VL-7 and PRB-1. (Table 2 & Figure 2).

**Table 2:** Effects of EMS on root length and shoot length in two varieties, VL-7 and PRB-1 of *F. esculentum*.

| TREATMENT | ROOT LENGTH (CM) |  |  |  | SHOOT LENGTH (CM) |  |  |  |
| --- | --- | --- | --- | --- | --- | --- | --- | --- |
| | $\bar{X}$ | S.D | S.E | C.V | $\bar{X}$ | S.D | S.E | C.V |
| VL-7 CONTROL | 3.03 | 0.55 | 0.31 | 0.18 | 1.83 | 0.76 | 0.44 | 0.41 |
| EMS (%) |  |  |  |  |  |  |  |  |
| 0.01 | 3.83 | 0.15 | 0.08 | 0.03 | 2.10 | 1.04 | 0.60 | 0.48 |
| 0.02 | 2.76 | 0.25 | 0.14 | 0.09 | 2.00 | 0.50 | 0.28 | 0.25 |
| 0.03 | 2.66 | 0.57 | 0.33 | 0.21 | 1.80 | 0.26 | 0.15 | 0.14 |
| 0.04 | 2.26 | 0.25 | 0.14 | 0.11 | 1.00 | 0.50 | 0.28 | 0.50 |
| CONTROL PRB-1 | 3.10 | 0.76 | 0.44 | 0.24 | 1.83 | 0.92 | 0.53 | 0.31 |
| EMS% |  |  |  |  |  |  |  |  |
| 0.01 | 3.50 | 0.50 | 0.28 | 0.14 | 2.10 | 1.04 | 0.60 | 0.56 |
| 0.02 | 2.93 | 0.92 | 0.53 | 0.31 | 2.10 | 0.52 | 0.30 | 0.25 |
| 0.03 | 2.83 | 0.76 | 0.44 | 0.26 | 1.83 | 1.04 | 0.60 | 0.56 |
| 0.04 | 2.46 | 0.15 | 0.08 | 0.06 | 1.00 | 0.50 | 0.28 | 0.50 |

**Fig. 2(a).**
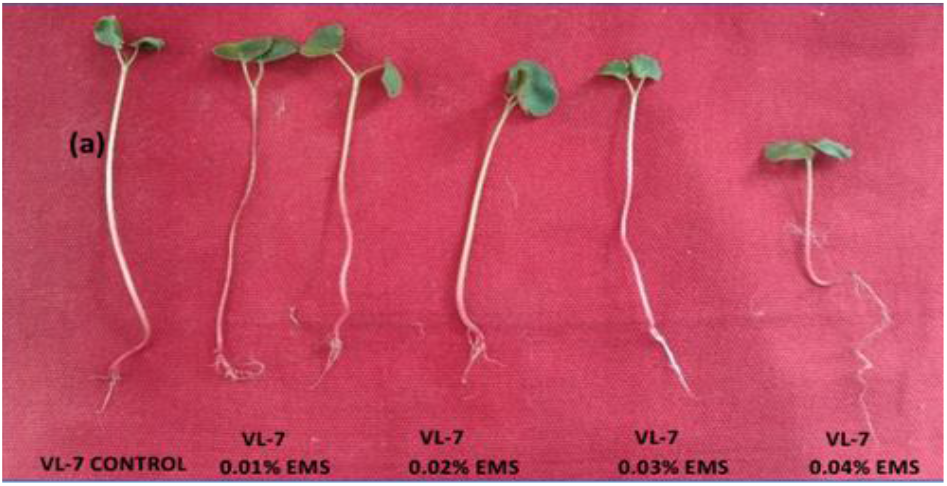
Buckwheat seed showing seedling height in control and EMS treatments (0.01% - 0.04%) in VL-7 variety.

**Fig.2 (b).**
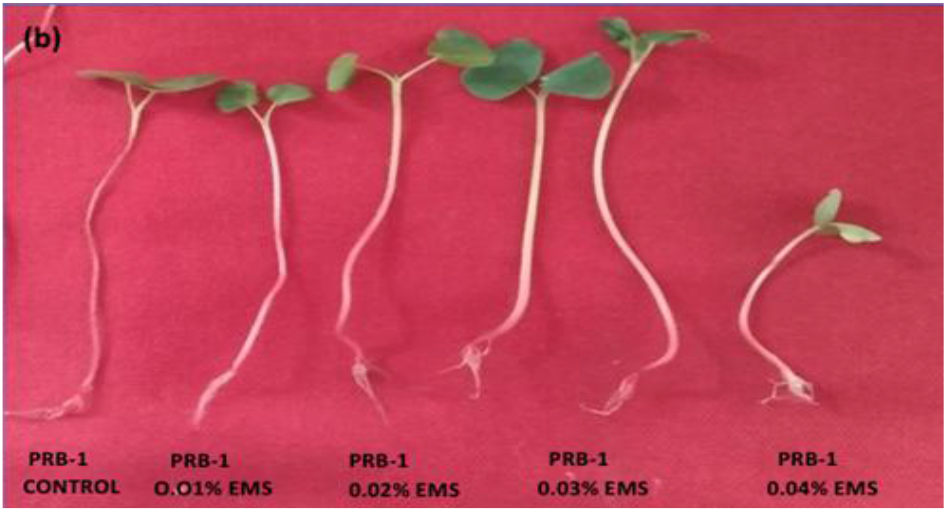
Buckwheat seed showing seedling height in control and EMS treatments (0.01% - 0.04%) in PRB-1 variety

#### Root length of seedlings

The root length in control was 6.40 cm and 6.88 cm in VL-7 and PRB-1, respectively. It decreased with increasing concentration of mutagen in PRB -1 from 7.50 cm to 4.90 cm and from 6.80 cm to 4.90 cm in VL-7. And the mean value for control was 3.03 and 3.10 in VL-7 and PRB-1, respectively, which decreased from 0.01% EMS to 0.04% EMS, from 3.83 cm to 2.26 cm in VL-7 and from 3.50 cm to 2.46 cm in PRB-1 (Table 2 & Figure 2).

#### Changes in the morphology of leaves

The shape of the leaves was normal in control of both varieties, but showed a huge variation in all the concentrations of mutagen in both varieties of *F. esculentum*. The variations observed were in the bifoliate leaves pattern, surface area, lamina morphology, degree and types of incision, and types of leaf bases, leaf apex, shape and size of the leaves, chlorotic leaves, high anthocyanin, elongated leaves, broad leaves, etc. (Figures 3 & 4).

**Fig. 3.**
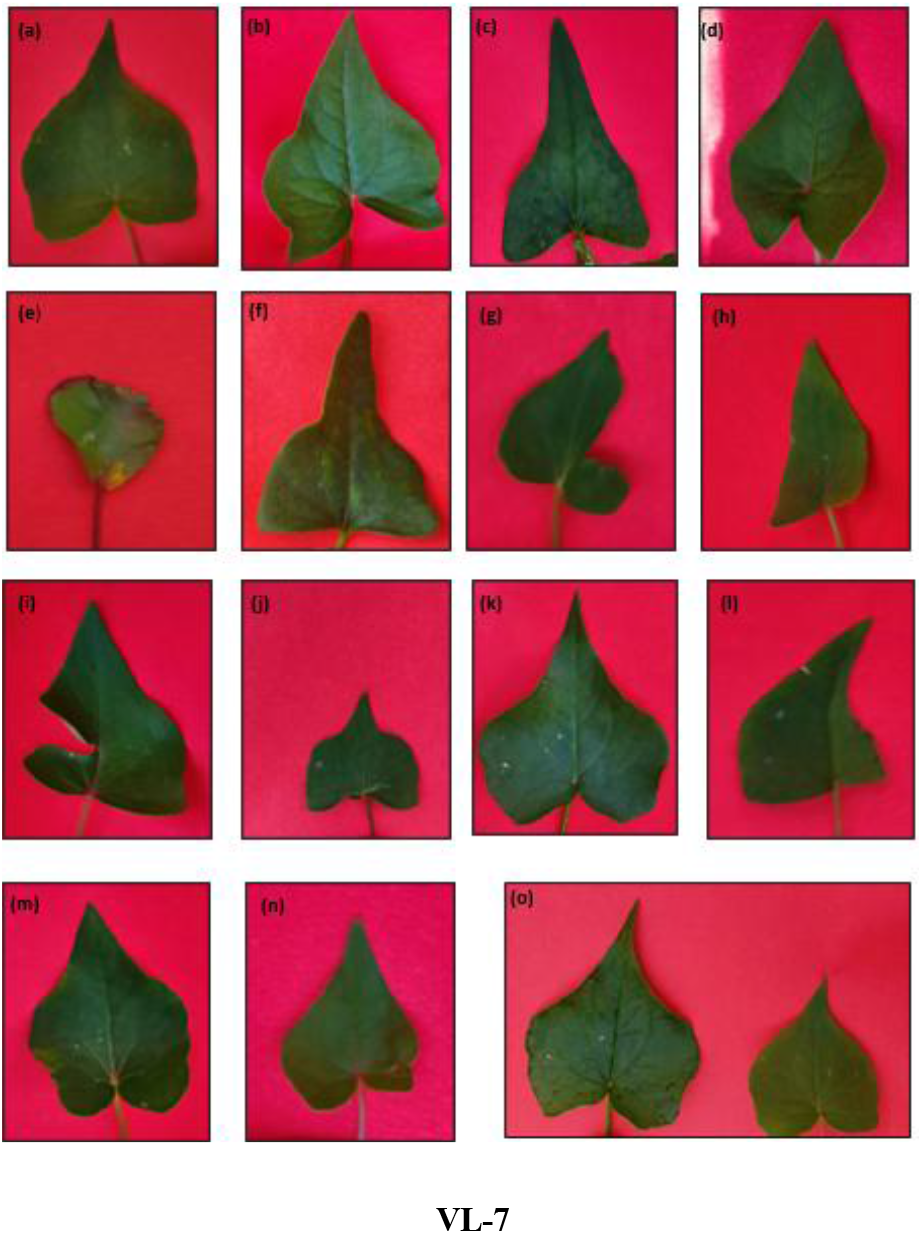
Morphological and chlorophyll abnormalities in leaves of VL-7 variety (a) Control leaf; (b) Oblique and sagittate base (0.01% EMS); (c) Elongated leaf (0.01% EMS); (d) High notch along with sagittate leaf base (0.01% EMS); (e) Tigrina leaf (0.01% EMS); (f) Truncate leaf base, notch is absent and aristate leaf apices (0.01% EMS); (g) Distorted leaf shape (0.01% EMS); (h) Oblique leaf base with no notch and elongated lamina (0.01% EMS); (i) Truncate leaf base and a large incision in leaf blade (0.01% EMS); (j) Sagittate base with incision, increased width in lamina along with an outgrowth and abruptly acuminate leaf apices (0.03% EMS); (k) Distorted leaf shape and truncate leaf base (0.01% EMS); (l) Caudate (0.01% EMS); (m) Irregular outlines of the blade, less pointed apex and more rounded (0.01% EMS); (n) Sagittate leaf base and leaf is broadly acute with small incision in lamina (0.01% EMS); (o) Comparison of reduction of leaf size in 0.03% EMS mutagenic treated leaf (right) with control (left).

#### Plant height and fertile branches

The average plant height in control of VL-7 was 27 cm, and in control of PRB-1 was 27.66 cm. In both the varieties, it first decreased in concentration of 0.01% EMS and then increased up to 0.04% EMS from 20.33 cm to 61.75 cm in VL-7 and 16.5 cm to 35.66 cm in PRB-1 (Table 3). Tall mutants with longer internodes and dwarf mutants with stunted growth and short internodes, thicker stems, were encountered with higher doses and lower doses of EMS, respectively, in both the varieties (Figures 5 & 6). In PRB-1, bushy mutants were observed in both tall and dwarf plants, having larger leaves and more flowers on fertile branches in the treatment of 0.03% and 0.01% EMS, respectively (Figure 6). Plants with a smaller number of fertile branches were observed in intermediate doses of EMS, whereas the plants with a greater number of fertile branches were observed in higher and lower doses of EMS in both varieties.

**Table 3:**
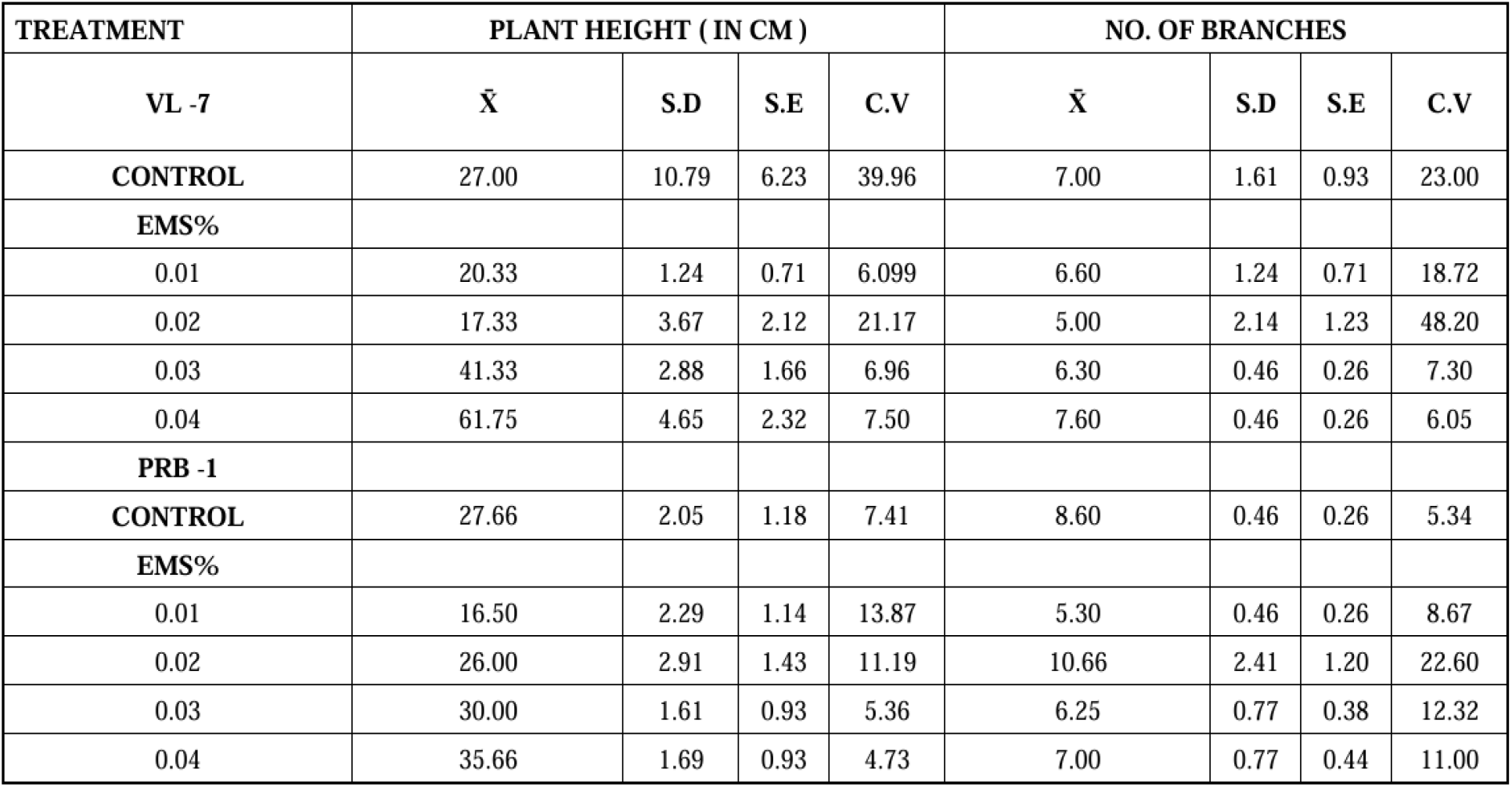
Effects of EMS on plant height and number of branches per plant in two varieties, VL -7 and PRB-1of *F. esculentum*.

| TREATMENT | PLANT HEIGHT ( IN CM ) |  |  |  | NO. OF BRANCHES |  |  |  |
| --- | --- | --- | --- | --- | --- | --- | --- | --- |
| VL -7 | $\bar{X}$ | S.D | S.E | C.V | $\bar{X}$ | S.D | S.E | C.V |
| CONTROL | 27.00 | 10.79 | 6.23 | 39.96 | 7.00 | 1.61 | 0.93 | 23.00 |
| EMS% |  |  |  |  |  |  |  |  |
| 0.01 | 20.33 | 1.24 | 0.71 | 6.099 | 6.60 | 1.24 | 0.71 | 18.72 |
| 0.02 | 17.33 | 3.67 | 2.12 | 21.17 | 5.00 | 2.14 | 1.23 | 48.20 |
| 0.03 | 41.33 | 2.88 | 1.66 | 6.96 | 6.30 | 0.46 | 0.26 | 7.30 |
| 0.04 | 61.75 | 4.65 | 2.32 | 7.50 | 7.60 | 0.46 | 0.26 | 6.05 |
| PRB -1 |  |  |  |  |  |  |  |  |
| CONTROL | 27.66 | 2.05 | 1.18 | 7.41 | 8.60 | 0.46 | 0.26 | 5.34 |
| EMS% |  |  |  |  |  |  |  |  |
| 0.01 | 16.50 | 2.29 | 1.14 | 13.87 | 5.30 | 0.46 | 0.26 | 8.67 |
| 0.02 | 26.00 | 2.91 | 1.43 | 11.19 | 10.66 | 2.41 | 1.20 | 22.60 |
| 0.03 | 30.00 | 1.61 | 0.93 | 5.36 | 6.25 | 0.77 | 0.38 | 12.32 |
| 0.04 | 35.66 | 1.69 | 0.93 | 4.73 | 7.00 | 0.77 | 0.44 | 11.00 |

**Fig. 4.**
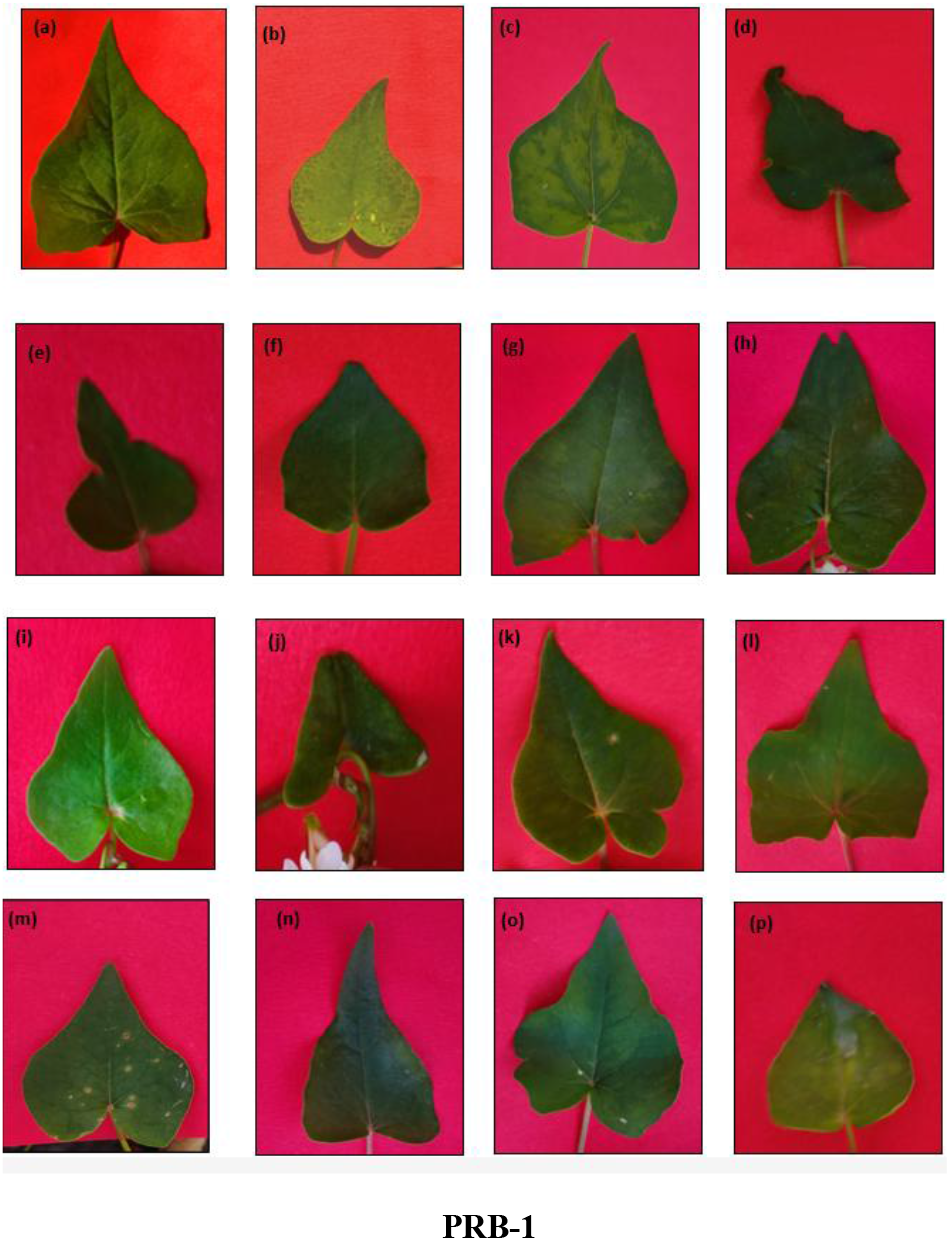
Morphological and chlorophyll abnormalities in leaves of PRB-1 variety (a) Control leaf; Variations in EMS treatment of PRB-1 (O.O2% EMS) from (b) to (l) : (b) Chlorina with dark green spots on the leaf; (c) Chlorosis with Irregular Mottling; (d) Distorted leaf shape with irregular boundaries and many incisions at the blade; (e) Truncate leaf base, aristate apex and the incision on both side of the lamina; (f) Heart shaped leaf; (g) A combination of oblique and sagittate type leaf base with incision; (h) Highly sagittate leaf base and the pointed apex is splitted in two leaf apices; (i) High notch, sagittate type leaf base and broadly acuminate; (j) Cotyledon type leaf shape; (k) Sagittate with incisions giving a round shaped structure at the base; (l) Truncate leaf base with incision in the middle at the notch and abruptly acuminate. (m) Red lesions on the leaf – microbial infection (0.04% EMS); (n) Elongated leaf and truncate leaf base (0.03% EMS); (o) Sagittate leaf base, incisions on one side of the leaf, and irregular in shape and aristate apex; (p) More rounded shape (0.04 % EMS).

**Fig. 5.**
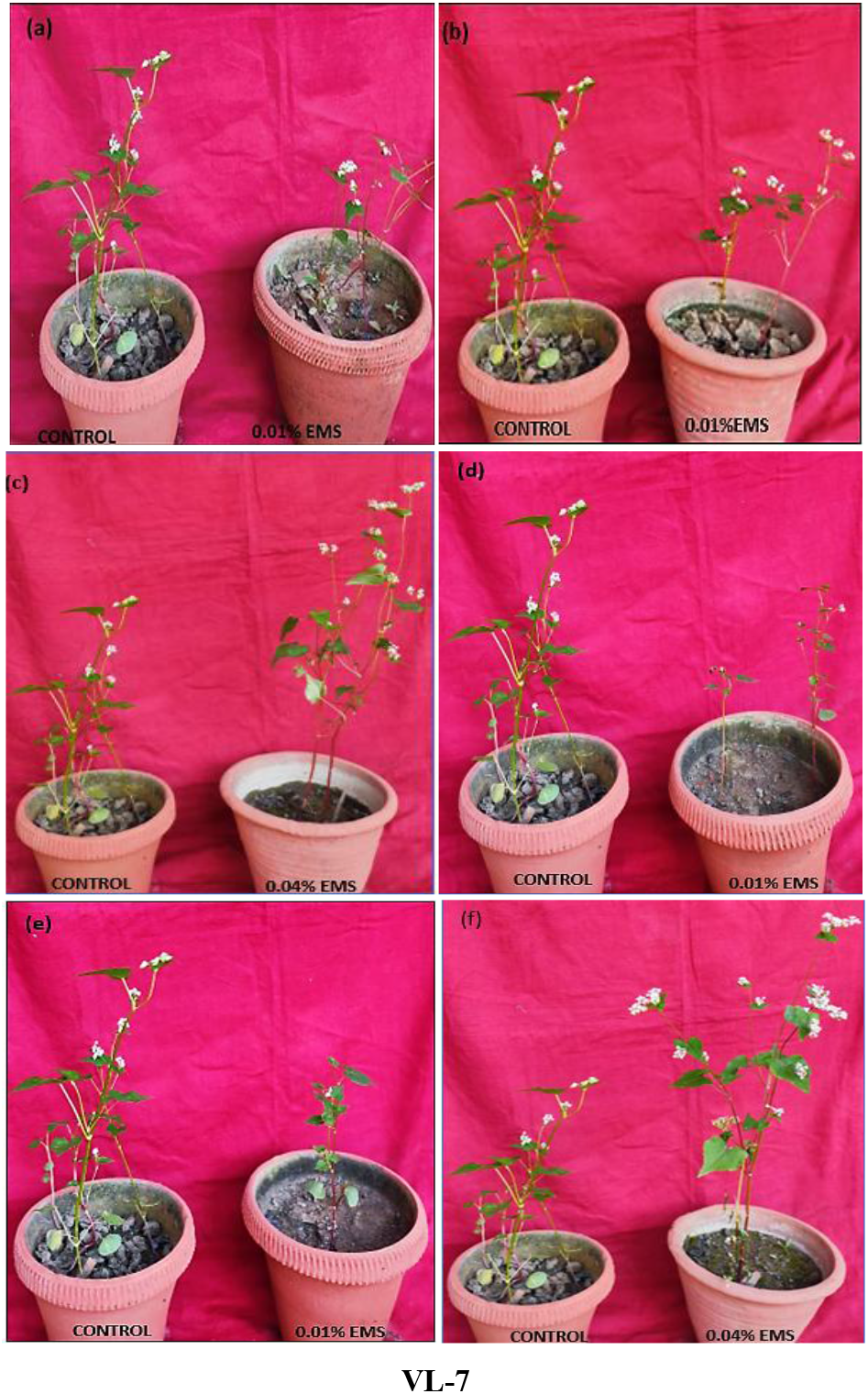
(a)-(b) Plant height in the treated plants (0.01% EMS) which are dwarf against the control set; (c) Plant height in the treated plants (0.04% EMS) which are tall against the control set; (d) - (e) Plant with a smaller number of fertile branches in the treated plants (0.01% EMS) against the control set; (f) Plant with a greater number of fertile branches in the treated plants (0.04% EMS) against the control set

**Fig. 6.**
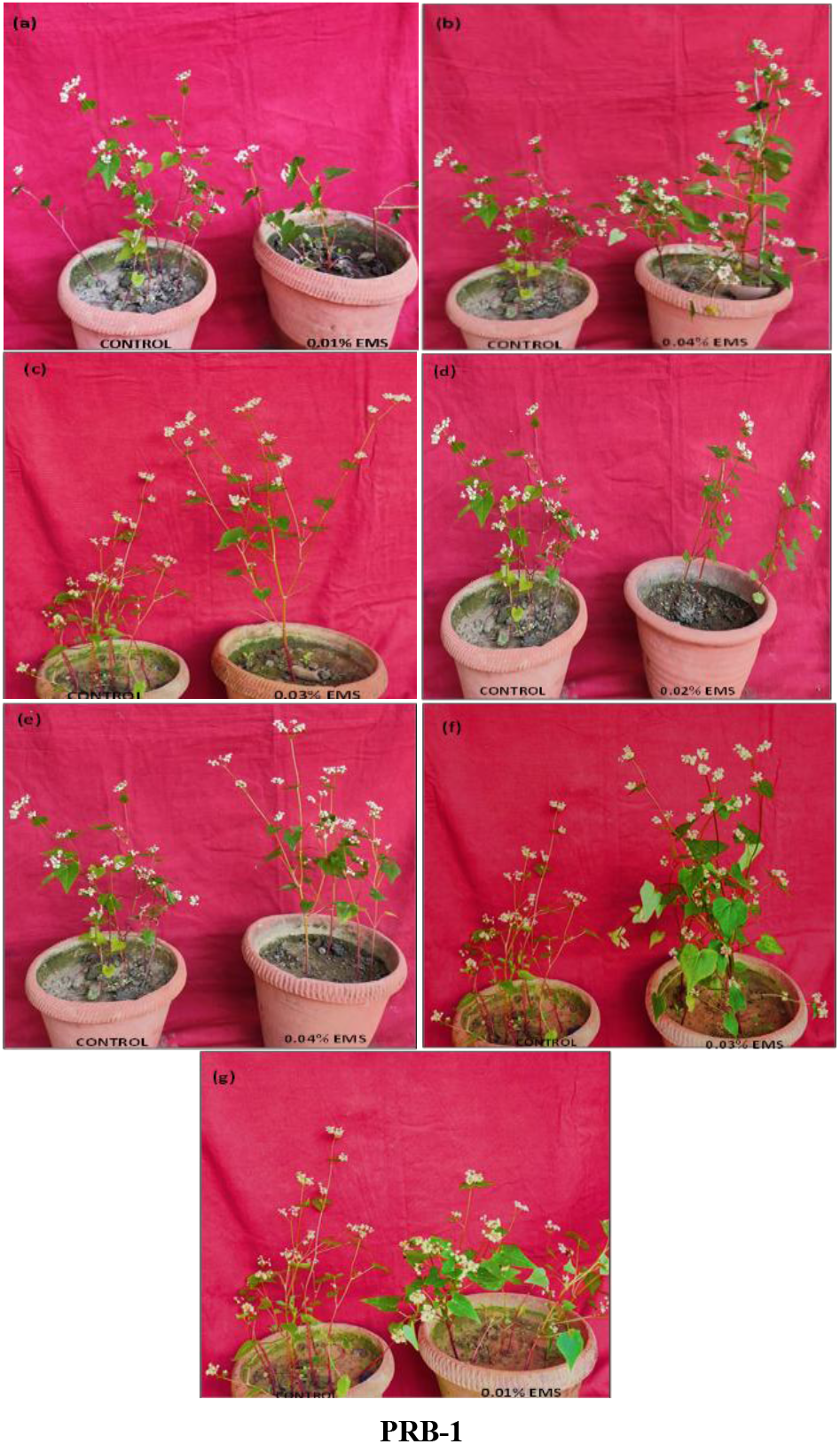
(a) Plant height in the treated plants (0.01% EMS) which are dwarf against the control set; (b) Plant height in the treated plants (0.04% EMS) which are tall against the control set; (c) Plant height in the treated plants (0.03% EMS) which are tall against the control set; (d) Plant with a smaller number of fertile branches in the treated plants (0.02% EMS) against the control set; (e) Plant with a greater number of fertile branches in the treated plants (0.04% EMS) against the control set; (f) A bushy variant that is tall, having larger leaves and more flowers on fertile branches (0.03% EMS) against the control set; (g) A bushy variant that is dwarf but have more and larger leaves and more flowers on fertile branches (0.01% EMS) side against the control set.

#### Number of branches

The average number of branches in control of VL-7 and PRB-1 was 7.00 to 8.60, respectively. It decreased in both the varieties, VL-7 and PRB-1, from 0.04% EMS to 0.01% EMS concentration as 7.60 to 6.60 and 7.00 to 5.30, respectively (Table 3).

#### Morphology and days to flowering

The average days to flowering in control of VL-7 was 62.00, and in control of PRB-1, it was 52.66. Then in VL-7 it decreased from 0.01% EMS to 0.04% EMS, from 55.33 to 51. In PRB-1, it decreased from 0.01% EMS to 0.04 % EMS, from 58.00 to 49.33 (Table 4). Some Hypopetalous mutant flowers were noticed in the VL-7 variety in a lower concentration of EMS treatment, where the number of petals reduced from five to four. Mutants with a smaller size of flower, different morphology of petals, and variations in anthers & stigma were also observed in lower doses of mutagen in both varieties (Figure 7).

**Table 4:** Effects of EMS on days to flowering and days to maturity in two varieties, VL -7 and PRB-1 of *F. esculentum*.

| TREATMENT | DAYS TO FLOWERING |  |  |  | DAYS TO MATURITY |  |  |  |
| --- | --- | --- | --- | --- | --- | --- | --- | --- |
| | $\bar{X}$ | S.D | S.E | C.V | $\bar{X}$ | S.D | S.E | C.V |
| VL-7 |  |  |  |  |  |  |  |  |
| CONTROL | 62.00 | 2.15 | 1.24 | 3.46 | 79.33 | 0.45 | 0.26 | 0.56 |
| EMS (%) |  |  |  |  |  |  |  |  |
| 0.01 | 55.33 | 2.05 | 1.18 | 3.70 | 77.33 | 1.69 | 0.97 | 2.18 |
| 0.02 | 60.33 | 1.24 | 0.71 | 2.05 | 75.50 | 0.50 | 0.35 | 0.66 |
| 0.03 | 52.00 | 0.81 | 0.46 | 1.55 | 72.00 | 1.22 | 0.61 | 1.69 |
| 0.04 | 51.00 | 1.41 | 0.81 | 2.76 | 71.25 | 0.82 | 0.41 | 1.15 |
| PRB-1 |  |  |  |  |  |  |  |  |
| CONTROL | 52.66 | 0.45 | 0.26 | 0.85 | 71.66 | 0.45 | 0.26 | 0.62 |
| EMS (%) |  |  |  |  |  |  |  |  |
| 0.01 | 58.00 | 0.81 | 0.46 | 1.39 | 75.66 | 0.99 | 0.57 | 1.30 |
| 0.02 | 56.00 | 1.87 | 0.93 | 3.30 | 71.75 | 0.82 | 0.47 | 1.14 |
| 0.03 | 50.75 | 0.86 | 0.215 | 1.69 | 70.00 | 0.70 | 0.40 | 1.00 |
| 0.04 | 49.33 | 0.93 | 0.53 | 1.88 | 68.35 | 0.45 | 0.26 | 0.65 |

**Fig. 7.**
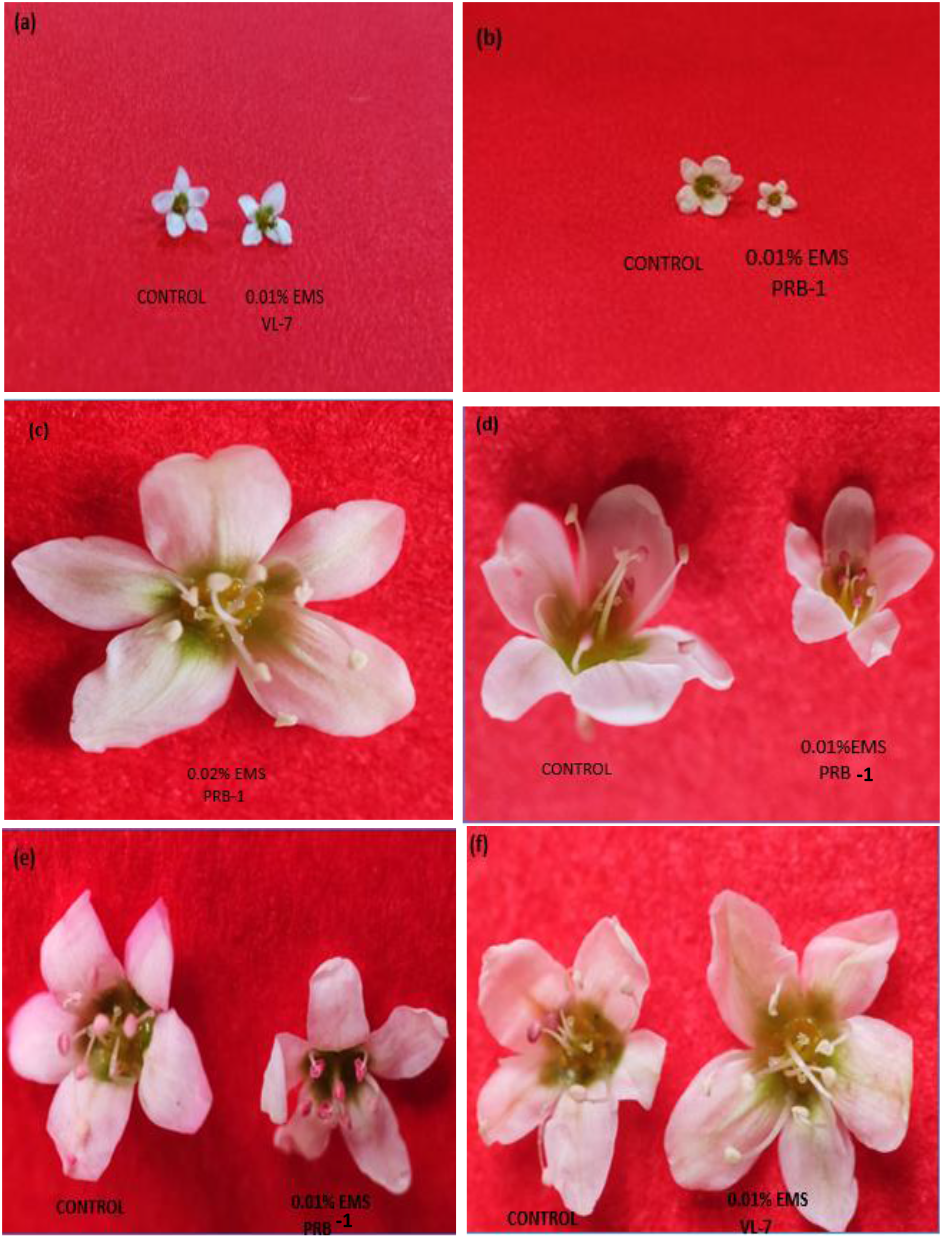
(a) Tetramerous flower in VL-7 mutagenic treatment 0.01% EMS against the pentamerous flower of control (Hypopetalous mutant); (b) Mutant with smaller flower size in 0.01% EMS of PRB-1 compared to the larger flower size of control; (c) Dimorphic flower mutant in 0.02% EMS PRB-1 having two types of petals, elongated and obovate in shape; (d) – (e) Mutants with short anthers in the flowers of mutagen treated 0.01% EMS of PRB1 against the long anthers of control (right); (f) Mutant Anthers and stigma completely white in 0.01% EMS of VL-7 and pink anthers in control.

#### Days to maturity

The average days to maturity in control of VL-7 were 79.33, and in PRB-1, it was 71.66. As the concentration increased from 0.01% EMS to 0.04% EMS, the average days to flowering decreased in both the varieties from 77.33 to 71.25 in VL-7 and 75.66 to 68.35 in PRB-1 (Table 4).

#### 100–Seeds weight (g)

The average seed weight in the control population of VL-7 and PRB-1 was 1.10 gm and 3.36 gm respectively. In VL-7 maximum weight was found in 0.04% EMS, which was 3.64 gm. In PRB-1, it first increased and then decreased from concentration 0.01% EMS to 0.04% EMS as 4.36 gm to 2.61 gm, respectively (Table 5)

**Table 5:** Effects of EMS on 100 seed weight and yield per plant in two varieties, VL - 7 and PRB -1 of *F. esculentum*.

| TREATMENT | 100 seed yield (gm) |  |  |  | yield per plant (gm) |  |  |  |
| --- | --- | --- | --- | --- | --- | --- | --- | --- |
| | $\bar{X}$ | S.D | S.E | C.V | $\bar{X}$ | S.D | S.E | C.V |
| VL-7 |  |  |  |  |  |  |  |  |
| CONTROL | 1.10 | 0.09 | 0.05 | 8.23 | 2.35 | 0.27 | 0.15 | 11.50 |
| EMS (%) |  |  |  |  |  |  |  |  |
| 0.01 | 3.32 | 0.10 | 0.06 | 2.96 | 3.37 | 0.23 | 0.13 | 6.78 |
| 0.02 | 1.82 | 0.01 | 0.01 | 0.66 | 2.06 | 0.05 | 0.02 | 2.42 |
| 0.03 | 1.50 | 0.03 | 0.02 | 2.06 | 2.59 | 0.24 | 0.14 | 9.20 |
| 0.04 | 3.64 | 0.36 | 0.21 | 9.92 | 6.15 | 0.05 | 0.03 | 0.66 |
| PRB-1 |  |  |  |  |  |  |  |  |
| CONTROL | 3.36 | 0.30 | 0.17 | 7.98 | 7.17 | 0.06 | 0.04 | 0.87 |
| EMS (%) |  |  |  |  |  |  |  |  |
| 0.01 | 4.36 | 0.34 | 0.20 | 7.91 | 5.82 | 0.01 | 0.01 | 0.13 |
| 0.02 | 3.24 | 0.36 | 0.21 | 11.06 | 3.72 | 0.18 | 0.11 | 4.95 |
| 0.03 | 3.23 | 0.34 | 0.197 | 10.57 | 4.45 | 0.23 | 0.13 | 5.09 |
| 0.04 | 2.61 | 0.05 | 0.03 | 1.73 | 4.41 | 0.36 | 0.21 | 8.25 |

#### Total seed yield per plant (g)

The average seed yield per plant in control of VL-7 was 2.35 g, and in control of PRB-1 it was 7.17 gm. Then in VL-7, it increased from 0.01% EMS 0.04% EMS from 3.37gm to 6.15gm respectively. In PRB-1, the maximum average yield per plant was found in the control, and then yield decreased with increasing concentration from 0.01% EMS to 0.04% EMS, as 5.82 g to 4.41gm respectively (Table 5).

#### Other abnormalities in mutated plants

Mutant plants having bifoliate leaflet-number and stunted growth, which later failed to survive, were observed in lower doses of EMS treatment of VL-7 and PRB-1 compared to the control set having trifoliate leaf pattern and normal growth. Bifoliate xantha, tigrina, and maculata mutants were also observed in lower doses of EMS treatment in both varieties. These were the mutants affected by the chlorophyll biosynthesis, yellowing of leaves, irregular patches on the leaf surface exhibiting stunted growth, and, eventually, mortality was observed. An increment in pigments, like high anthocyanin production, with an intensified pink pigmentation, was evident along the venation pattern. Cotyledonary leaves showing pinkish spots and other mutants with irregular pinkish patches were also recorded in the hyperanthocyanic mutant population of VL-7, 0.01% EMS and 0.04% EMS treatments. Also, the hyperveination mutant plants with 0.02% EMS treatment in VL-7 were observed with the number or density of veins per leaf area higher than the normal. (Figure 8).

**Fig. 8.**
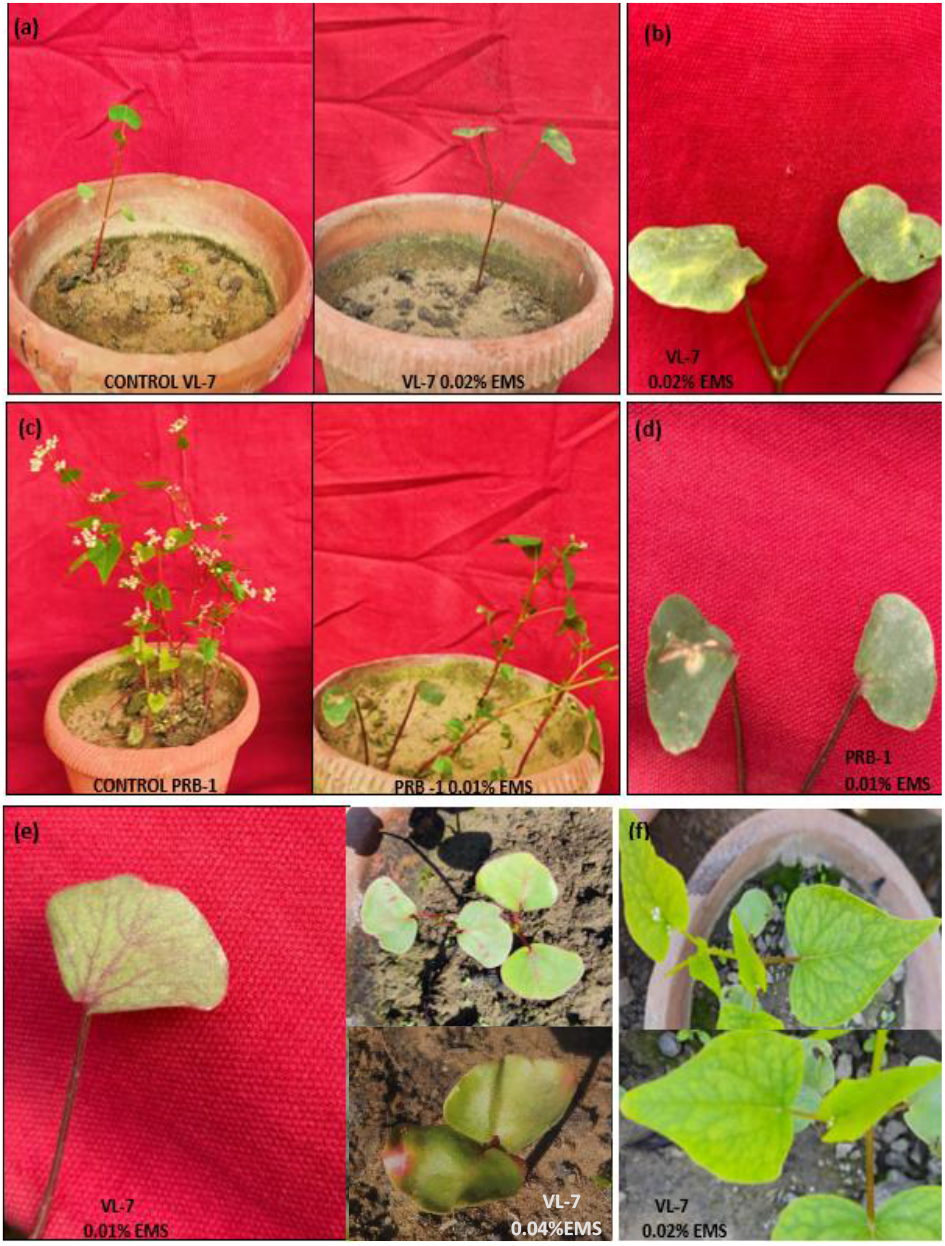
(a) Mutant plant having a bifoliate leaf pattern and stunted growth against the control set having a trifoliate leaf pattern and normal growth; (b) Xantha–Tigrina Mutant; (c) Mutant plant having a bifoliate leaf and stunted growth against the control set having a trifoliate leaf pattern and normal growth; (d) Bifoliate maculata mutant; (e) Hyperanthocyanic mutants; (f) Hyperveination mutant

#### Cytological abnormalities

Cytological analysis of EMS treated population of *F. esculentum* resulted in the formation of abnormal meiotic products. Compared to the chromosomes of control set with normal meiotic behaviour, VL-7 showed unequal chromosomes number at Telophase l and diagonal Anaphase l in 0.04% EMS and 0.02% EMS treatments respectively. Also, in PRB-1 unequal Telophase was recorded in 0.01% EMS treatment (Figure 9).

**Fig. 9.**
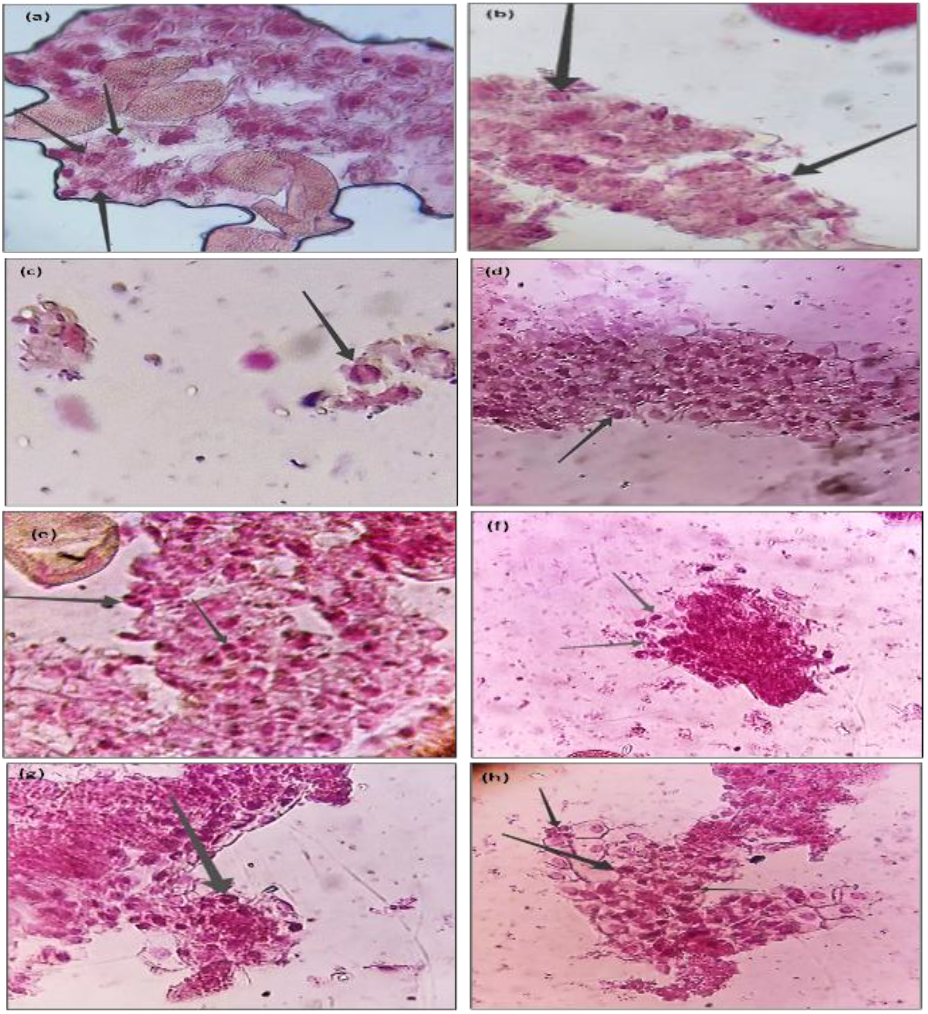
(a)Metaphase ll and Anaphase l (control); (b) Telophase l and Anaphase l (VL-7 0.01% EMS); (c) Telophase l (PRB-1 0.02% EMS); (d) Unequal chromosomes number at Telophase l (VL-7 0.04% EMS); (e) Diagonal Anaphase l and normal Telophase l (VL-7 0.02 % EMS); (f) Telophase l (PRB -1 0.03% EMS); (g) Unequal Telophase (PRB-1 0.01% EMS); (h) Metaphase l, Telophase l and Telophase ll (PRB-1 0.04% EMS)

### Physiological studies

#### Chlorophyll content (mg/ml)

The chlorophyll content in control of the VL-7 was 0.262 mg/ml, and in control of PRB-1 it was 0.113 mg/ml. As moving from concentration 0.01% EMS to 0.04% EMS, first it decreased, then increased from 0.132 mg/ml to 0.212 mg/ml in VL-7, but in PRB 1 it decreased from 0.215 mg/ml to 0.146 mg/ml in concentration 0.01% EMS to 0.04% EMS (Table 6).

**Table 6:** Effects of EMS on chlorophyll and carotenoid content in two varieties, VL -7 and PRB -1 of *F. esculentum*.

| TREATMENT | CHLOROPHYLL (mg /ml) | CAROTENOID (mg / ml) |
| --- | --- | --- |
| VL -7 |  |  |
| CONTROL | 0.262 | 0.053 |
| EMS (%) |  |  |
| 0.01 | 0.132 | 0.033 |
| 0.02 | 0.206 | 0.046 |
| 0.03 | 0.204 | 0.047 |
| 0.04 | 0.212 | 0.048 |
| PRB -1 |  |  |
| CONTROL | 0.113 | 0.039 |
| EMS (%) |  |  |
| 0.01 | 0.215 | 0.047 |
| 0.02 | 0.197 | 0.048 |
| 0.03 | 0.217 | 0.047 |
| 0.04 | 0.146 | 0.043 |

#### Carotenoid content (mg/ml)

Carotenoid content in the control set of VL-7 was 0.053 mg/ml, and in PRB-1 it was 0.039 mg/ml. In VL-7, it increased from 0.033 to 0.048mg/ml from 0.01% EMS to 0.04% EMS. And in PRB-1, it first increased from 0.01% EMS and decreased up to 0.04%EMS as 0.047 to 0.043 mg/ml (Table 6)

## Discussion

Enhancement of frequency and spectrum of mutations in a predictable manner, thereby achieving desired plant characteristics, is an important goal of mutation research. The present study has proved fruitful in inducing a wide range of morphological, biological, and cytological mutations. Such mutants might be either a result of pleiotropic effects of mutated genes or chromosomal aberrations or gene mutations (Khan *et al*., 2011). The tall, dwarf, bushy, etc. mutants may not be useful for direct commercial cultivation; they, however, may be used in hybridization to transfer their useful traits to other high-yielding varieties of buckwheat. Though it is not easy to eliminate the negative traits of the pleiotropic spectrum from the positive ones, the pleiotropic pattern of the mutant gene can, however, be altered to some extent by transferring it into a specific genotypic background (Sidorova, 1981). Various workers reported that mutant types like tall, dwarf, bushy, and leaf were monogenic and recessive in behavior (Jana, 1963; Sharma & Sharma, 1981; Reddy & Gupta, 1988; Satyanarayana *et al*., 1989; Singh *et al*., 1999; Qin *et al*., 2008; Talukdar, 2009). However, Konzak *et al*. (1969) in wheat and Shakoor *et al*. (1978) in triticale reported semi-dwarf character to be polygenic in nature. Morphological mutants, isolated in M_1_ generation, differed not only in the two varieties of buckwheat but also within the variety in different mutagenic treatments, suggesting that the varieties responded differently to the dose and type of mutagens employed. Relative differences in the mutability of genes for different traits have been observed, as some of the mutant types, like mutations in leaf and height, appeared with higher frequencies than the others in some mutagens. The more frequent induction of certain mutation types by a particular mutagen may be attributed to the fact that the genes for these traits are probably more responsive to different mutagens with different modes of action. Tall mutants, as observed in our experiment, were also reported earlier by Solanki *et al*. (2004) in lentil and Kumar *et al*. (2009) in black gram. Jana (1963) observed tall mutants in X-ray treatments in black gram and reported that tallness was due to longer internodes. The longer internodes may be due to the high gibberellic acid content. The tall mutant found in higher concentrations of EMS (0.04% and 0.03%) in our experiment had thinner and longer stems, which may be because they invested a significant proportion of their biomass towards the stems to increase length rather than towards lateral expansion of stem diameter, resulting in weak stems that lodge easily and decreased the yield. The dwarf mutant that was observed in lower concentrations of EMS (0.01% and 0.02%) may be suitable for selection as they have higher breaking force and lower lodging index that may provide a high-yielding variety. The dwarf character may be due to the lower GA_4_ and GA_7_ content and higher Jasmonic acid, Salicylic acid, and brassinosteroids. The positive correlation of stem girth, which is related to the prevention of stem lodging at the maturity stage, is an important desirable character in buckwheat that was observed in dwarf mutants. Bifoliate mutants like tigrina, xantha, and maculata exhibiting stunted growth and eventually mortality were observed. Bifoliate mutation affects leaflet initiation/organogenesis, usually controlled by KNOX, PHAN/ARP, or auxin-patterning genes. These chlorophyll mutants were observed to survive only a limited number of days or weeks, because they couldn’t sustain sufficient photosynthesis (Raina *et al*., 2022). These pigment mutations result from genetic changes (mutations) in genes involved in chlorophyll synthesis or chloroplast biogenesis — sometimes affecting nuclear genes, plastid genes, or genes regulating pigment production (Yu *et al*., 2007). Such mutants help in the study of genes involved in chlorophyll synthesis, chloroplast development, light response, and other basic plant biology. Several hyperanthocyanic mutants were observed with higher-than-normal levels of anthocyanins, which may be due to alterations in transcription factors, structural genes in the flavonoid pathway, and stress-response or signaling pathways that regulate anthocyanin biosynthesis. Many early-maturing mutants were isolated at various mutagen treatments in both varieties. It was observed that the plants that were dwarf, having small, shorter, and fewer stems and branches was seen with early flowering. The shorter stem, fewer and shorter branches, and fewer internodes lead to the allocation of resources towards supporting structures and early flowering, i.e., condensed but complete life cycle to maintain population persistence. Earliness is one of the reliable characteristics of mutation experiments. Lateness is a less desired mutant character (Gottschalk & Wolff, 2012). Increase in branching obtained may be due to quick cell division and elongation, and phytohormones or nucleic acids biosynthesis. The mutations in flowers, like the shifting of pentamerous flowers toward the tetramerous, may be due to the variation in the ABC model. The number of flowers per plant and the number of seeds per inflorescence are the main components contributing to yield. Various types of leaf mutants, such as narrow, broad, fused, bi-, and multifoliate, were recorded in both varieties. Considerable variability was also observed among the leaves in terms of surface area, lamina morphology, degree and types of incision, and types of leaf bases. The plant with narrow leaf character gives the larger seed that is useful for a better groat yield. Such leaf changes were attributed to chromosomal breakage, disturbed auxin synthesis, disruption of mineral metabolism, and accumulation of free amino acids (Gunkel & Sparrow, 1961). Morphological mutants were assayed for nitrate reductase activity (NRA), total chlorophyll and carotenoid contents, which differed from the control. The role of NRA and chlorophyll contents in promoting the growth and enriching the nutritional quality of the crops through induced mutagenesis has been reported in chickpea (Barshile *et al*., 2009; Barshile & Apparoa, 2009). A mutation in seed weight is another desirable trait for evaluating plant yield, which was increased in medium doses of EMS. In the present experiment, in the PRB-1 variety, the isolated dwarf, bushy, early maturing, and bold-seeded mutants exhibited the higher values for total carotenoid contents with respect to the controls, depicting their role in stress physiology. In higher plants, carotenoids protect the photosynthetic apparatus from excess of photons and oxidative stress, which are generated under stress (Siefermann-Harms, 1987). The induced mutants were often associated with cytological abnormalities. Meiosis is one of the most important genetic events occurring in meiocytes of the organism, consisting of highly balanced biochemical, cytogenetic, physiological, and phenotypic events leading to chromosome reduction, gene rearrangements, and gamete formation (Goyal *et al*., 2019). Cytology of mutants revealed normal meiotic divisions and the abnormal diagonal anaphase and unequal telophase. Narrow leaf mutant was also observed with a moderate dose of EMS, resulting in meiotic abnormalities and unequal separation in telophase l may be due to the laggard chromosome formation and the diagonal anaphase I, or may be due to the disruption in spindle fibre and disrupt polarity. One of the research articles suggests that the unequal separation of chromosomes was due to early or delayed separation of bivalents and multivalents in the mutated forms of chromosomes, and it may result in the formation of aneuploid gametes (Zeerak, 1992).

## Summary

Mutants are essential basic materials in genetic research and play a critical role in genetic research and quality improvement. The establishment of a mutant library is also important for accelerating crop breeding and broadening plant genetic resources. Researchers have created crop mutants and bred new plant varieties. However, whether these mutant materials can aid in molecular-assisted breeding still needs to be determined, and determining whether their mutation is stable requires ongoing planting of the next generation to observe their mutation stability. In addition, analysis of the causes of mutations from a genomic perspective has become an essential direction of functional genomics research. Due to the self-pollination mode, buckwheat possesses a narrow genetic base, which requires substantial research to broaden its genetic variability. One possible method to improve diversity and create high-yielding lines is to introduce mutations into the buckwheat genome. Since then, only a few mutagenesis studies on buckwheat have been conducted, and only a few mutant populations have been created, with just a few cultivars released. As a result, additional mutagenesis researches are required to produce genetic diversity that will aid in the development of superior cultivars of desirable features. This study investigated the mutagenic effects of an important chemical mutagen, Ethyl Methane Sulphonate (EMS), on *F. esculentum*. Buckwheat seeds were treated with four different concentrations of EMS (0.01%, 0.02%, 0.03%, 0.04%). The main objective was to enhance and to improve the diversity and create high-yielding lines. So, the study mainly focused on – Biological damage in M_1_ generation and inducing variability in morphological traits. It was observed that the induction and extent of genetic variability using EMS in buckwheat seed depend on the mutagen doses. The treatment was found to be appropriate for breeding objectives aimed at productivity, considering the lower biological damage and higher rate of variation in quantitative traits, along with unique morphological variations. The significant findings are summarized below-

1. The seed germination shows an increasing trend with the decreasing concentration of mutagen.
2. The inhibition decreased with decreasing concentration of the mutagen.
3. The plant survival increased with decreasing concentration of the mutagen doses.
4. Studies on various quantitative parameters revealed the general effectiveness of intermediate doses and stimulatory effectiveness of lower and higher concentrations in M_1_ generation.
5. The average height of mature plants first reduced in a lower concentration of mutagen, then increased up to a higher concentration in the M_1_ generation.
6. Considerable variability of leaf mutants and floral mutants was studied in different doses of mutagens.
7. Several bifoliate and chlorophyll mutants like tigrina, maculata, xantha, viridis, and pigment mutant plants like hyperanthocyanic, along with hyperveination mutant were studied in lower doses of treatments.
8. Cytological analysis shows metaphase I, telophase I, telophase II, and anaphase I with abnormalities like diagonal anaphase I and unequal telophase in different concentrations of mutagen, and also the pollen fertility was affected.
9. Various other characters, such as the average number of branches per plant, seeds per plant, and yield per plant, height of the plant, bushiness of the plant, fertile branches per plant, variations in leaf, and morphological variations in flowers, were studied in M_1_ generation. All these characters first decreased and then increased with increasing concentrations of mutagen, and it was found 0.01% EMS and 0.04% EMS were better yielding concentration amongst all.

The findings of the present mutation breeding study can be explored further in M_2_ generation to understand the genetic response and permanent mutations in *F. esculentum* towards different doses of EMS, which will definitely provide a resource for setting mutation breeding protocols and superior varieties.

## Acknowledgement

The authors are thankful to Aligarh Muslim University, Aligarh, India, for institutional facilities that enabled the successful completion of this work and Prof. Samiullah Khan and Dr Sana Chaudhary (Assistant Professor) for their guidance and support.

## Author Contribution

Ayesha Badar: conceptualization, methodology, validation, formal analysis, investigation, data curation, writing—original draft, writing—review and editing, visualization, project administration; Iram Siddique: resources; Hakeem Mubeen and Iram Siddique: supervision.

